# A Chemogenetic Platform for Spatio-temporal Control of β-arrestin Translocation and Signaling at G protein-Coupled Receptors

**DOI:** 10.1101/251769

**Authors:** Yusuke Gotoh, Patrick M. Giguere, David E. Nichols, Bryan L. Roth

## Abstract

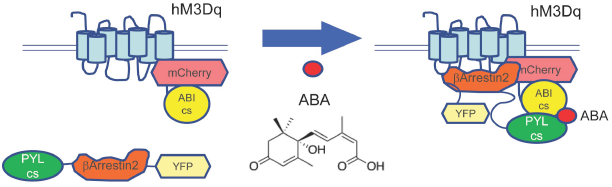

Although ligand-activated GPCRs induce both G-protein and β-arrestin dependent signaling, gaining precise spatio-temporal control of β-arrestin signaling has proven elusive. Here we describe a platform for specifically activating β-arrestin-dependent signaling *in situ*. The platform, which we have dubbed “GA-PAIR” (<u>G</u>PCR/β-<u>A</u>rrestin –<u>P</u>lant protein and <u>A</u>bscisic acid <u>I</u>nduced <u>R</u>ecruitment), can be controlled by the inert phytochemical S-(+)-abscisic acid (ABA). ABA induces interaction between ABI1 (ABA Insensitive 1) and PYL1 (Pyrabactin Resistance (PYR) 1-Like), two plant proteins with no mammalian counterparts. We fused ABI to the engineered human muscarinic M3 G protein-coupled receptor (hM3Dq) and PYL1 to β-arrestin2. Addition of ABA induced rapid and nearly complete translocation of the PYL-β-arrestin fusion protein and, importantly, induced both ERK and Akt signaling. Photo-uncaging a new photo-caged ABA analogue allowed us to gain relatively precise spatio-temporal control over β-arrestin translocation. Because GA-PAIR facilitates the exclusive activation of endogenous β-arrestin signaling pathways in the absence of a GPCR ligand or G protein, the GA-PAIR system will facilitate deconvoluting GPCR signaling *in situ.*

## INTRODUCTION

G protein-coupled receptors (GPCRs) are the largest superfamily of signaling molecules in the human genome^1^ and, as such, play key roles for transducing extracellular stimuli into intracellular signaling. GPCRs are also key regulators of many diseases and prospective targets for drugs. Indeed, more than 30% of the drugs used in the clinic target GPCRs.^2^ Therefore, elucidating the role of GPCRs is very important for understanding normal biology, disease pathology and therapeutic drug discovery. GPCRs are generally activated via binding of extracellular ligands, which trigger conformational changes resulting in activation of their associated hetero-trimetric G proteins.^1^ G protein activation engages multiple cellular signaling pathways, for example, cAMP induction, cAMP reduction, and/or Ca^2+^ release, typically referred to as “G-protein signaling” or “canonical GPCR signaling”. Following activation, GPCRs are rapidly phosphorylated by G protein–coupled receptor kinases (GRKs)^3,4,5^ and in most instances, the phosphorylated GPCRs interact with β-arrestin family proteins (e.g. β-arrestin1 and/or β-arrestin2).^5^ This GPCR-arrestin interaction was originally described as a way to terminate GPCR signaling via interference with G protein coupling and by promoting GPCR internalization by targeting the GPCRs to clathrin-coated pits.^6,7,8^

During the past decade, however, it has become increasingly clear that β-arrestins can serve as adaptor proteins to transduce signals to multiple effector pathways, including ERK (Extracellular signal-Regulated Kinase) and various other signaling pathways.^5,9,10,11,12,13^ This β-arrestin signaling is now known as a major non-canonical GPCR signaling pathway.^1^ Most natural ligands and chemical agonists of GPCRs are thought to activate both the canonical and non-canonical GPCR pathways. Recently, however, it has been verified that there are some agonists that can disproportionately activate only one of the two signaling branches, a phenomenon usually referred as functional selectivity or biased signaling.^14,15,16^ Although some progress has been made using G-protein or β-arrestin biased agonists to delineate the physiological functions of these signaling pathways,^17,18^ because no ligand displays absolute bias their utility has been limited. The ideal system would afford the exclusive and generic activation of β-arrestin translocation in the absence of any G-protein activation. For G protein dependent signaling analysis, we have previously described a platform known as DREADD (Designer Receptor Exclusively Activated by Designer Drug) in which directed molecular evolution in yeast was used to engineer a receptor that is solely activated by a pharmacologically inert, drug-like small molecule such as clozapine-N-oxide (CNO).^19,20^ To date, Gq, Gi, and Gs coupled-DREADDs (hM3Dq, hM4Di, and GsD respectively) have been created and extensively validated^21^. These DREADDs allow for chemogenetic activation of various G protein dependent signaling pathways in essentially any cellular context^21^.

Currently there are several methods available preferentially to activate β-arrestin dependent signaling. For example, biased agonists can selectively activate β-arrestin and they have been used to reveal many important observations regarding β-arrestin dependent signaling.^13,22,16^ A β-arrestin-biased DREADD Rq (R165L) also has been engineered that can selectively activate β-arrestin dependent signaling without perturbing G protein dependent signaling.^23^ Rq (R165L) represents a powerful proof-of-concept for selective activation. It should be noted, however, that more than 1 μM of CNO is required to activate β-arrestin dependent signaling using Rq(R165L). This high concentration of CNO likely results in off-target receptor activation and undesirable additional signaling events. Additionally, the low potency of CNO is not likely to be particularly useful *in vivo*, and because it is CNO-sensitive, the Rq(R165L) DREADD cannot be used in combination with G-protein-biased DREADDs^21^ limiting this approach.

Another method to generate isolated β-arrestin dependent signaling was developed in 2004 by Terrillon et al.^24^ wherein β-arrestin2 is recruited to the membrane using the FKBP-FRB dimerization system. This approach, however, has been limited by the instability, toxicity, and off-target effects of AP21967, the rapamycin analogue used to activate the FKBP-FRB system^25^. In addition, this method is complicated by the use of FKBP and FRB fusion proteins, which exist abundantly in mammalian systems. As a result, endogenous mammalian FKBP and cyclophilin compete for binding of FKBP and FRB fusion proteins.

Recently, it was reported that the phytohormone S-(+)-abscisic acid (ABA)^26^ can be used to induce the interaction between two plant proteins, ABI1 (ABA Insensitive 1) and PYL1 (Pyrabactin Resistance (PYR) 1-Like). The interacting complementary surfaces (CSs) of PYL1 (PYLcs, amino acids 33 to 209) and ABI1 (ABIcs, amino acids 126 to 423) confer ABA-induced proximity to proteins fused to these fragments.^26^ Although the ABA signaling pathway mediates stress responses and developmental decisions in plants, ABA is nontoxic in humans^27^ and indeed, is present in low amounts in our diet.^27^

Here we provide a system wherein the interaction between hM3Dq and β-arrestin is controlled by ABA-induced dimerization of ABIcs and PYLcs. We call this newly developed system “GA-PAIR” for (<u>G</u>PCR/β-<u>A</u>rrestin-<u>P</u>lant proteins and <u>A</u>bscisic acid <u>I</u>nduced <u>R</u>ecruitment) system. GA-PAIR represents an excellent tool for elucidating the physiological role of β-arrestin signaling *in vitro* and, potentially, *in vivo*.

## RESULTS AND DISCUSSIONS

### ABA induces rapid membrane translocation of a β-arrestin fusion construct

In order to induce membrane translocation of β-arrestin and subsequent β-arrestin dependent signaling without GPCR ligands we designed two fusion proteins; hM3Dq-mCherry-ABIcs and PYLcs-β-arrestin2-YFP (**Fig. 1a**). Using an automated calcium-mobilization assay, we found that fusing the ABIcs protein to the C-terminal of hM3Dq-mCherry did not significantly affect Gq signaling induced by CNO (**Fig. 1b**), indicating the fused receptor to be fully functional.^19^ Using live-cell confocal imaging acquisition in HEK293T cells co-expressing both fusion proteins, we found that hM3Dq-mCherry-ABIcs was located at the plasma membrane as expected, while PYLcs-β-arrestin2-YFP was detected throughout the cytoplasm prior to ABA stimulation (**Fig. 1c:** Time = 0). After exposure to 200 μM ABA, PYLcs-β-arrestin2-YFP rapidly translocated to the membrane, and this process was almost complete after 2 minutes of ABA exposure (**Fig. 1c**). Different combinations and orientations of the fusion proteins were tested and the hM3Dq-mCherry-ABIcs and PYLcs-β-arrestin2-YFP were found to be the most efficient pairing (not shown).

**Figure 1.**
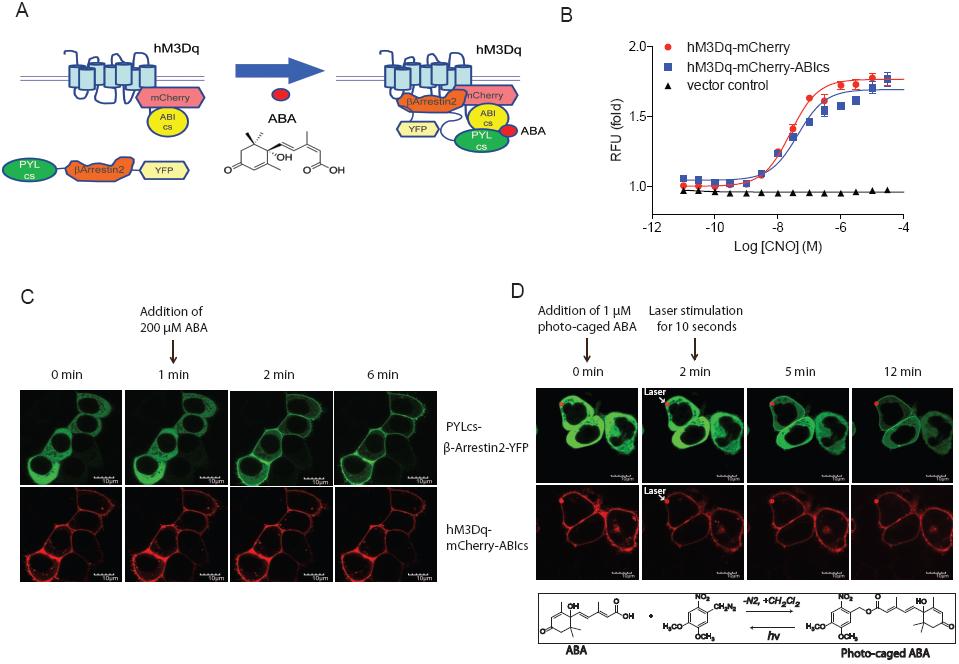
Membrane translocation of PYLcs-β-arrestin2-YFP induced by ABA stimulation. (a) Schematic representation of ABA-induced translocation of PYLcs-β-arrestin2-YFP to hM3Dq-mCherry-ABIcs. (b) FLIPR assay for validation of Gq activity of the fusion receptors. HEK293T cells transfected with hM3Dq-mCherry (red line), hM3Dq-mCherry-ABIcs (blue line), or vector control (black line) were stimulated with CNO, and the Gq signal was measured by FLIPR^Tetra^. (c). Time course images of HEK 293T cells co-transfected with hM3Dq-mCherry-ABIcs and PYLcs-β-arrestin2-YFP. ABA was added to the cells at time = 1 minute. (d-upper panel) Time course images of HEK 293T cells co-transfected with hM3Dq-mCherry-ABIcs and PYLcs-β-arrestin2-YFP. The cells were incubated with 1 μM photo-caged ABA for 2 min and were stimulated with the 405 nm laser for 10 seconds at time = 2 minutes in the red circle as indicated. Scale bar represents 10 μm. (d-bottom panel) Photo-caged ABA (2-nitro-4,5-dimethoxybenzyl ester of (±)-abscisic acid) synthesis. Prepared in a single step from abscisic acid and the diazo compound obtained from the MnO_2_ oxidation of 2-nitro-4,5-dimethoxybenzaldehyde hydrazide.

In order to control the spatio-temporal translocation of the β-arrestin, we next generated a photo-caged version of ABA (2-Nitro-4,5-dimethoxybenzyl ester of Abscisic acid) (**Fig. 1d, bottom panel)**. Cells co-expressing both hM3Dq-mCherry-ABIcs and PYLcs-β-arrestin2-YFP were incubated with 1 μM photo-caged ABA for 2 minutes, and then stimulated for 10 seconds using a 405 nm laser illumination. Prior to laser illumination, PYLcs-β-arrestin2-YFP was located throughout the cytoplasm (**Fig. 1d:** Time = 2 minutes). Following laser illumination at a specific focal point (red circle), the PYLcs-β-arrestin2-YFP robustly translocated in the illuminated cell (**Fig. 1d:** Time = 5 minutes) and weakly in a neighboring cell (**Fig. 1d:** Time = 12 minutes). The data demonstrate that it is possible to control β-arrestin2 translocation within a specific cell using photo-caged ABA and limited laser illumination to gain a more precise spatio-temporal control.

### Optimization of β-arrestin signaling

β-arrestin-mediated signaling via GPCR recruitment can transduce signals to multiple effector pathways, including ERK and various transcriptional events.^9,10,11,12,13^ The Serum Response Element luciferase reporter assay (SRE-luc) is commonly used as a sensitive approach to detect ERK-dependent signaling pathway activity^28,29^ and we used it to measure ERK-dependent signaling induced by ABA in our engineered platform.

HEK293T cells were then co-transfected with hM3Dq-ABIcs and PYLcs-β-arrestin2-YFP or β-arrestin2-YFP (e.g. lacking the PYLcs domain) as a negative control. CNO stimulation, which activates G-protein and β-arrestin dependent signaling, resulted—as expected—in a robust activation of SRE-luc with both β-arrestin fusion proteins (**Fig. 2a**). Consequently, ABA exposure, which solely induces β-arrestin dependent translocation, resulted in luciferase activity only in the cells expressing hM3Dq-ABIcs and PYLcs-β-arrestin2-YFP (**Fig. 2b**) while no luciferase activity in control cells expressing hM3Dq-ABIcs and β-arrestin2-YFP was observed (**Fig.2b**). These results demonstrate that β-arrestin2 translocation alone is sufficient to activate signaling as measured by SRE-luc activity.

**Figure 2.**
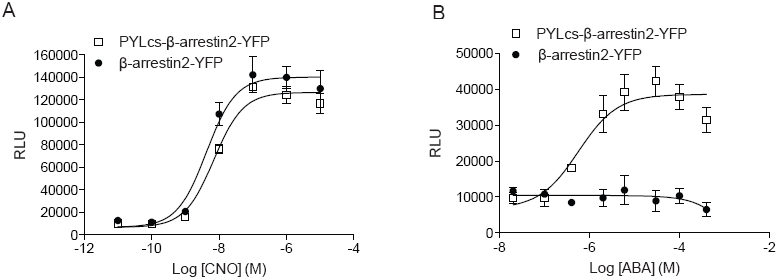
β-arrestin2-dependent signaling induced by the GA-PAIR system. (a and b). HEK293T cells were co-transfected with SRE-Luc, hM3Dq-ABIcs, and PYLcs-β-arrestin2-YFP (open square) or the control β-arrestin2-YFP (closed circle) and cells were stimulated with CNO (a) or ABA (b). After 4 h incubation, the luciferase activity was measured using the Bright-Glo assay system.

In order to optimize β-arrestin2 dependent signaling in GA-PAIR system, we modified each construct using insertional and targeted mutagenesis. First, we inserted the C-terminal fragment of human arginine vasopressin receptor 2 (AVPR2) (V2 tail) between hM3Dq and ABIcs. The V2 tail insertion was expected to enhance the activation of β-arrestin2 dependent signaling.^30^ HEK293T cells were transfected with hM3Dq-ABIcs and PYLcs-β-arrestin2-YFP or with hM3Dq-V2 tail-ABIcs and PYLcs-β-arrestin2-YFP. As shown in **Fig. 3b**, the luciferase activity induced by ABA stimulation was higher in the hM3Dq-V2 tail-ABIcs expressing cells than the hM3Dq-ABIcs expressing cells. This result indicates that the insertion of V2 tail between hM3Dq and ABIcs enhances β-arrestin2 dependent signaling as previously reported.^30^

**Figure 3.**
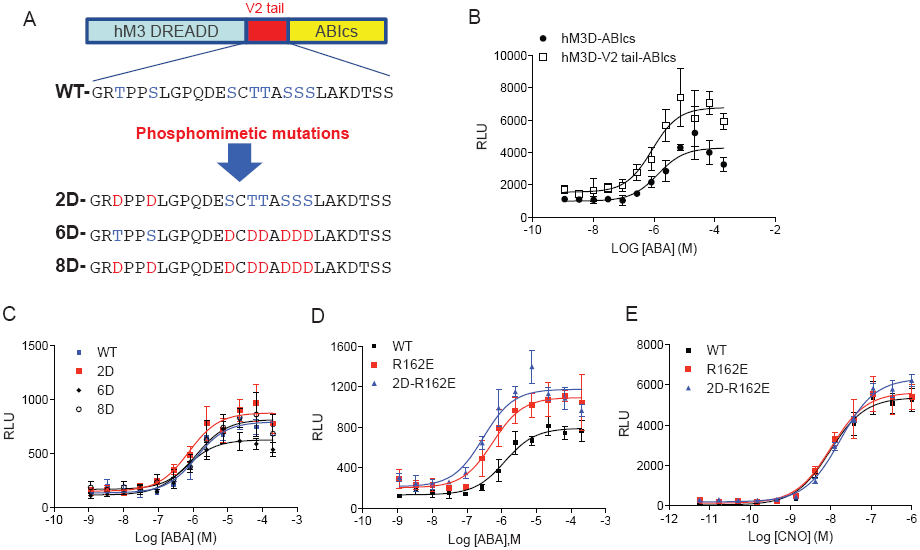
Optimization of the GA-PAIR system for β-arrestin2-dependent signaling. (a) Schematic representation of the V2 tail amino acid sequence and corresponding phosphomimetic mutations inserted between the C-terminal end of the receptor and the fusion ABlcs moiety. (b) HEK293T cells were co-transfected with SRE-Luc, PYLcs-β-arrestin2-YFP, and hM3Dq-ABIcs (closed circle), or hM3Dq-V2 tail-ABIcs (open square), and cells were stimulated with ABA. (c) HEK293T cells were co-transfected with SRE-Luc, PYLcs-β-arrestin2-YFP, and hM3Dq fusion proteins as follows; hM3Dq-ABIcs (blue line), hM3Dq-V2 tail (2D)-ABIcs (red line), hM3Dq-V2 tail, (6D)-ABIcs (black line diamond), or hM3Dq-V2 tail (8D)-ABIcs (black line open circle). (d) HEK293T cells were co-transfected with SRE-Luc, hM3Dq-V2 tail-ABIcs (WT), and PYLcs-β-arrestin2-YFP (WT) (black line), hM3Dq-V2 tail-ABIcs (WT) and PYLcs-β-arrestin2 (R162E)-YFP (red line), or hM3Dq-V2 tail (2D)-ABIcs and PYLcs-β-arrestin2 (R162E)-YFP (blue line). Cells were stimulated with ABA (c and d) or CNO (e) for 4 h and the luciferase activity was measured using the Bright-Glo assay system.

There are five serine and three threonine residues in the V2 tail sequence and its phosphorylation at serine and threonine residues by G protein-coupled receptor kinases (GRKs) induces activation and further high affinity β-arrestin association with activated GPCRs.^5^ To enhance β-arrestin2 recruitment and activation in our system in a GRK independent manner (i.e. no G-protein mediated activation), we mutated these residues to aspartic acid (D), a potential phosphomimetic of serine and threonine (**Fig. 3a**). These phosphomimetic mutations did not reduce Gq or SRE-luc activity when stimulated with CNO (**Supplementary Fig. 1a and Supplementary Fig. 1b**). The 2D mutant slightly enhanced SRE-luc activity following ABA treatment, however, whereas the 6D mutant showed a decrease, and the 8D mutant had no effect (**Fig. 3c**).

Several mutants of β-arrestin2 have been reported in the literature to modulate β-arrestin-receptor complex dynamics. For example, the R170E^31,32,33^ was found to interact with nonphosphorylated receptor, whereas the 3A (I386A, V387A, F388A)^33,34^ mutant mimics the active state of β-arrestin (receptor-bound conformation). We engineered these mutations into our PYLcs-β-arrestin2-YFP construct to determine which mutation(s) might enhance β-arrestin2 dependent signaling and no change was observed for the R170E and the 3A mutants (**Supplementary Fig. 2a and 2b**).

In 2013, Shukla et al.^35^ reported the crystal structure of β-arrestin1 bound to a V2 vasopressin receptor phosphopeptide, revealing for the first time the structure of the active conformation. This work revealed important structural change occurring in β-arrestin following interaction with a phosphorylated receptor peptide. Most of the reported changes occur in the receptor-binding cleft of the N domain and especially in the three distinctive finger loops, middle loop, and lariat loop. Several point mutations were generated in the cleft that is believed to be the key structural feature that regulates GPCR interactions with arrestins. One point mutation, R162E, was found to regulate β-arrestin2 functions and was tested in our platform for its ability to promote ERK signaling. As shown in **Figure 3d**, the R162E mutant appreciably enhances the SRE-luc activity in the presence of ABA (Emax: 789 (WT) vs 1093 (R162E); EC50: (1236 nM (WT) vs 551 nM (R162E)), but not CNO **(Figure 3e)**. The combination of the R162E mutation and the 2D phosphomimetic results in a synergistic increase in SRE reporter activation induced by ABA **(Figure 3d)** (Emax: 789 (WT) vs 1175 (2D-R162E); EC50: (1236 nM (WT) vs 293 nM (2D-R162E)). These results demonstrate that the R162E mutation is sufficient to enhance β-arrestin2 dependent signaling following membrane translocation.

β-arrestin2 dependent signaling has been demonstrated to activate multiple pathways in many cellular backgrounds,^13^ with ERK and Akt being common downstream effectors.^9,13,36,37^ To determine if the GA-PAIR system stimulates similar pathways we quantified their activation using antibodies directed against phospho-ERK, phospho-Akt, and phospho-PRAS40 (a substrates of Akt^38^) via Western blotting. For these studies, cells were transfected with hM3Dq-V2 tail (2D)-ABIcs and PYLcs-β-arrestin2 (R162E)-YFP and stimulated with ABA or CNO. As shown in **Figure 4a**, receptor activation with the full agonist CNO resulted in a 5-fold increase in ERK phosphorylation after 5 minutes. By contrast, β-arrestin2 translocation induced by ABA also resulted in a 1.5-fold increase in ERK phosphorylation after 5 minutes. These phospho-ERK measurements correlate with the results from our SRE-luc assay presented above. Additionally, β-arrestin2 translocation via ABA results in a 1.2-fold increase in Akt phosphorylation after 5 minutes (**Fig. 4b**). Furthermore β-arrestin2 translocation via ABA results in a 1.5-fold increase in PRAS40 phosphorylation after 5 minutes, and this signal is sustained for 30 minutes post-stimulation (**Fig. 4c**). These results demonstrate that β-arrestin2 dependent signaling induced by ABA activates the ERK and Akt pathways in the absence of GPCR ligand or Gq activation (**Fig. 5**).

**Figure 4.**
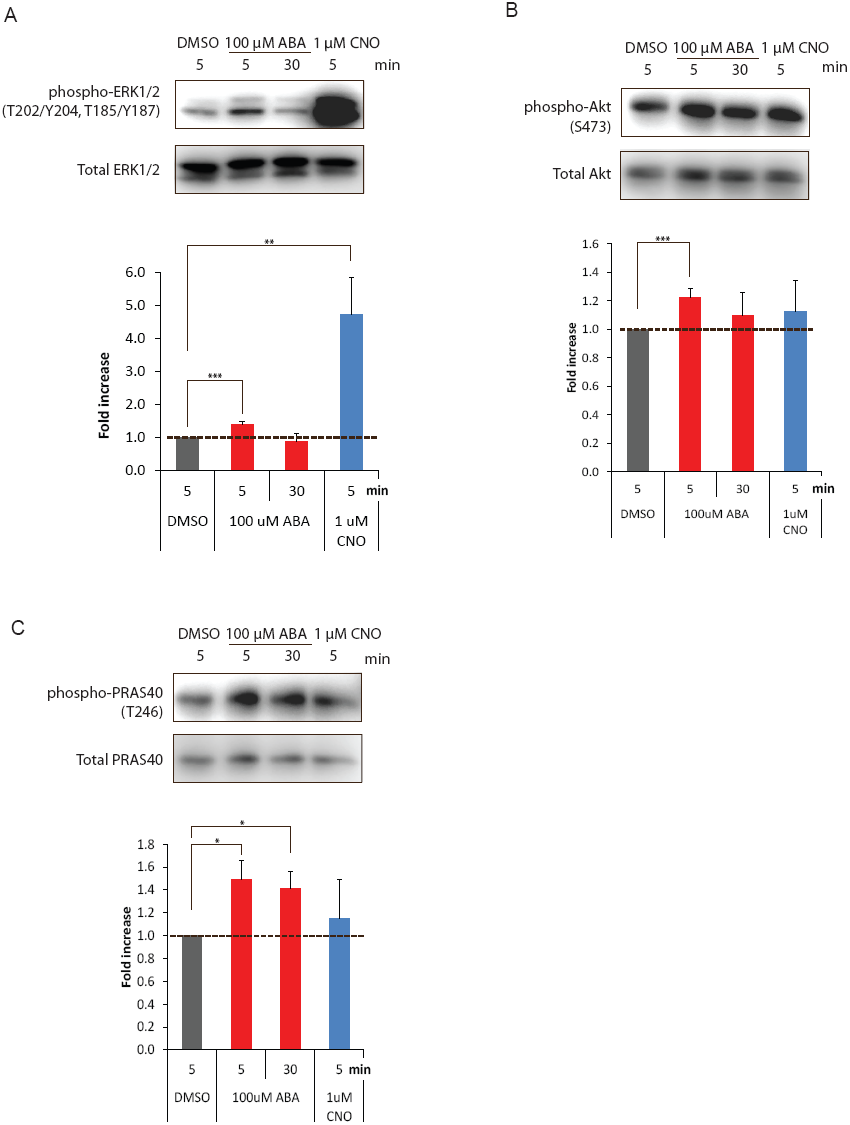
Quantitative analysis of ABA or CNO induced phosphorylation of (a) ERK, (b) Akt, and (c) PRAS40 by Western blotting assay. HEK293 cells transfected with hM3Dq-V2 tail (2D)-ABIcs and PYLcs-β-arrestin2 (R162E)-YFP were stimulated with 100 μM ABA for 5 or 30 min, or 1 μM CNO for 5 min. Western blotting assays were performed using whole cell lysate. Representative phosphoprotein and total protein immunoblotting images are shown above a bar graph depicting the mean of the fold increase of DMSO control ± standard deviation (SD). Band intensities were quantified using a GEL LOGIC 2200 IMAGING System (Kodak). Total observed phosphorylated signal for each protein was normalized to its respective total protein level. Each experiment was performed at least three times. Statistical analysis was done using Welch’s t-test. \**P* < 0.05, \*\**P* < 0.01, \*\*\**P* < 0.001.

**Figure 5.**
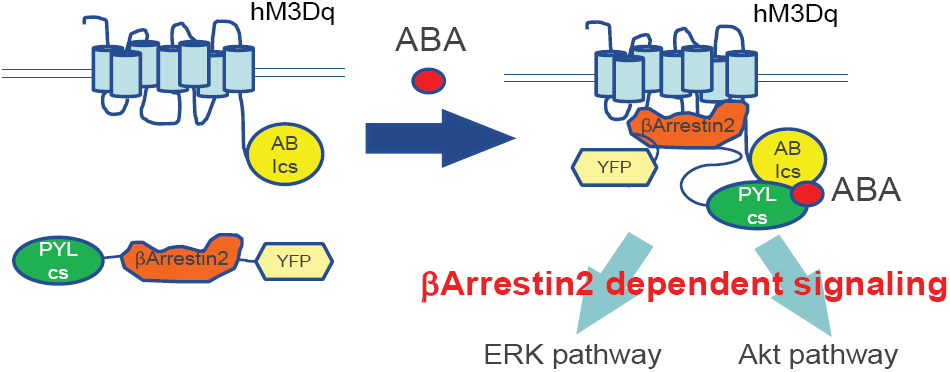
Schematic representation of ABA stimulation-induced β-arrestin2 dependent signaling. When cells expressing hM3Dq-V2 tail (2D)-ABIcs and PYLcs-β-arrestin2 (R162E)-YFP are stimulated with ABA, PYLcs-β-arrestin2 (R162E)-YFP translocates to the membrane and complexes with hM3Dq-V2 tail (2D)-ABIcs. The complex then induces β-arrestin2 dependent signaling, which involves the ERK and Akt pathways.

Here we describe the development and validation of a highly efficacious and apparently non-toxic system for the controlled translocation and activation of β-arrestin at an engineered GPCR. Through application of this system we have made three key discoveries. First, introduction of the V2 tail and the β-arrestin2 (R162E) mutation enhance β-arrestin2 dependent signaling. Second, β-arrestin2 signaling is sufficient to produce sustained ERK and Akt pathway activation in the absence of GPCR ligands or Gq activation. Finally, we were able to refine the method allowing for the more precise spatio-temporal control of β-arrestin2 signaling in a single cell using laser-activation of a newly designed photo-caged ABA.

hM3Dq was used in our system to act as an internal control, allowing us to monitor the effects of our fusion constructs on Gq activation and to afford control of canonical signaling in the absence of competing endogenous receptor. Also, the demonstration of the utility of the GA-PAIR system allows for the potential multiplexed use of our system sequentially to control canonical signaling using low doses of CNO and β-arrestin2 signaling via ABA. Although not yet attempted, the ultimate introduction of this system to experimental animals will allow us to distinguish comprehensively between the consequences of G protein and β-arrestin dependent signaling. Moreover, because we can easily substitute β-arrestin1 for β-arrestin2, the system could be used to interrogate the functional differences between β-arrestin1 and β-arrestin2 dependent signaling.

In conclusion, we have developed the GA-PAIR system, a controllable β-arrestin signaling system that operates independently of GPCR activation. Control of this system was further expanded using photo-caged ABA and laser stimulation specifically to target subcellular localization of β-arrestin. Through application of this system we demonstrate that β-arrestin alone is sufficient to activate both the ERK and Akt signaling pathways. This newly developed GA-PAIR system will represent a formidable platform for elucidating the physiological role(s) of β-arrestin signaling in vitro and in vivo.

## MATERIALS AND METHODS

### Chemical reagents, antibodies, and plasmids

S-(+)-abscisic acid (ABA) (Cat# 10073) was purchased from Cayman Chemical. Protease inhibitor cocktail (Cat# 11 873 580 001) was purchased from Roche. Phosphatase inhibitors cocktail 2 and 3 (Cat# P5726 and P0044, respectively) were purchased from Sigma. Transfection reagent TransIT-2020 (Cat# MIR5400) was purchased from Mirus. Antibodies for ERK1/2 (Cat# 9102), phospho-ERK1/2 (Thr202/Tyr204) (Cat# 9101), Akt (Cat# 4691), phospho-Akt (Ser473) (Cat# 4060), PRAS40 (Cat# 2610), and phospho-PRAS40 (Thr246) (Cat# 13175), were purchased from Cell Signaling. cDNAs of hM3Dq, β-arrestin2, ABIcs, PYLcs, mCherry and YFP were artificially synthesized by Blue Heron Biotech. These cDNAs were subcloned into a pcDNA3.1 vector. The pSRE-Luc reporter vector was purchased from Promega.

### Cell culture and transfection

HEK293 cells and HEK293T cells were grown in Dulbecco’s modified Eagle’s medium (Invitrogen) supplemented with 10% fetal bovine serum and 1× Penicillin/Streptomycin (Invitrogen) in humidified 5 % CO2 at 37 °C. Cells were transfected by lipofection using TransIT-2020 according to the manufacturer’s protocol.

### Translocation assay

HEK293T cells were co-transfected with the indicated ABIcs constructs and PYLcs constructs. After 24 h incubation, cells were harvested, suspended in 0.5% dialyzed FBS/DMEM, and plated in poly-L-lysine coated glass bottom 35 mm dishes (P35G-1.5-14-C, MatTek). After overnight incubation, the medium was changed to an observation buffer (1x Hank’s Balanced Salt Solution (HBSS), 20 mM HEPES pH 7.4, and 0.1% BSA) and cells were stimulated with ABA or photo-caged ABA, and live-imaged with a FV1000 confocal microscope (Olympus). In the case of laser activation of photo-caged ABA, the 405 nm laser was used under the control of the confocal microscope and the laser power was 50%.

### SRE-Luciferase reporter gene assay

HEK293T cells were co-transfected with the indicated ABIcs constructs, PYLcs constructs and pSRE-Luc vector. After 24 h incubation, the cells were harvested, suspended in 0.5% dialyzed FBS/DMEM, and plated at 20000 cells/well in poly-L-lysine coated 384-well white clear bottom plates. After overnight incubation, cells were stimulated with ABA or CNO, and luciferase activity was measured using the Bright-Glo Luciferase Reporter assay system (Promega) according to the manufacturer’s protocol.

### Western Blotting

The HEK293 cells transiently overexpressing the indicated ABIcs constructs and PYLcs constructs were serum starved overnight and treated with and without 200 μM ABA or 1 μM CNO for the indicated time at 37 °C. After treatment, cells were washed twice with ice-cold PBS and lysed for 30 min in the lysis buffer (50 mM Tris-HCl pH7.5, 150 mM NaCl, 5 mM EDTA, 1 mM DTT, 0.5% Triton X-100, protease inhibitor cocktail and phosphatase inhibitor cocktail 2/3). After centrifugation, cleared cell lysates were separated by SDS-PAGE and transferred to nitrocellulose membranes for Western blotting. Protein levels were detected by specific antibodies. Chemi-luminescence detection was performed using Super Signal West Pico reagent (Pierce). Band intensities were quantified by using the GEL LOGIC 2200 IMAGING System (Kodak). The phosphorylation level of each protein was normalized by the amount of each total protein. Each experiment was performed at least three times. For stripping the membrane, One Minute Advance Western Blot Stripping Buffer (GM Biosciences) was used.

### FLIPR assay

HEK293T cells were transfected with hM3Dq fusion protein. After 24 h, the cells were harvested, suspended in 0.5% dialyzed FBS/DMEM, and plated at 20000 cells/well in poly-L-lysine coated black 384-well clear bottom plates. After overnight incubation, medium was removed and the cells were loaded with 20 μL/well of 1x Fluo4 Direct Calcium dye (Invitrogen). Plates were incubated for 60 min at 37 °C, followed by 10 minute incubation at room temperature, and then loaded in the FLIPR^TETRA^. The FLIPR was programmed to take 10 readings (1 read per second), first as a baseline before addition of 10 μL of 3x CNO solution. The fluorescence intensity was recorded for 2 minutes after CNO addition. We measured maximal fluorescence intensity readings within one minute after CNO addition and calculated the Relative Fluorescence Units (RFU) and fold activity. For more details, see our PDSP (NIMH Psychoactive Drug Screening Program) protocol (http://pdsp.med.unc.edu/pdspw/function.php).

### Photo-caged (+)-ABA (2-Nitro-4,5-dimethoxybenzyl ester of Abscisic acid) synthesis

Commercial 2-nitro-4,5-dimethoxybenzaldehyde was obtained from Sigma and recrystallized once from EtOH. The aldehyde was converted to its hydrazide, and then to the diazo compound following the procedure described by Wootton and Trentham.^39^ A solution of 90.1 mg (0.4 mmol) of 2-nitro-4,5-dimethoxybenzaldehyde hydrazide was dissolved in 5 mL of dry CH_2_Cl_2_, followed by addition of 620 mg of activated MnO_2_ (Sigma). The reaction was carried out under reduced light, stirred at room temp for 45 minutes, then gravity filtered through Celite into a glass scintillation vial containing 105.7 mg (0.4 mmol) of dry (+)-abscisic acid with a magnetic stirring bar. The Celite pad was rinsed into the abscisic acid reaction with a further 7 mL of dichloromethane. The stirring diazonium solution was a dark red color. Vigorous gas evolution was initially observed, with loss of the dark red color. The resulting light orange reaction was placed in the cold room overnight to provide a nearly colorless solution. The solvent was then removed by rotary evaporator under reduced light to afford a pale yellow oil. This oil was dissolved in 2-3 mL of dry diethyl ether and then diluted with hexanes just to the cloud point. Storage in the cold room overnight, protected from light, led to the formation of crystals, which were collected by suction filtration, washed with hexane, and air-dried to afford 105 mg of light pink crystals. The mother liquor was concentrated and a second crop of 34 mg was obtained; total yield 139 mg (76%). The crystals had mp 96-106 °C (dec). The NMR was consistent with the expected structure and the cims showed M+2 at 461.2 m/z, with the expected λ_max_ at 350 nm^40^. No effort was made to determine photolysis kinetics, however, lc-ms analysis of an ethanol solution of the product following 350 nM UV irradiation for 5 min generated an eluting peak with the same retention time as authentic abscisic acid and a corresponding mass ion for abscisic acid at 247.2 m/z (M+H-18). The abscisic acid peak was absent from the sample prior to UV irradiation.

## SUPPLEMENTARY FIGURE LEGENDS

**Supplementary Figure 1.**
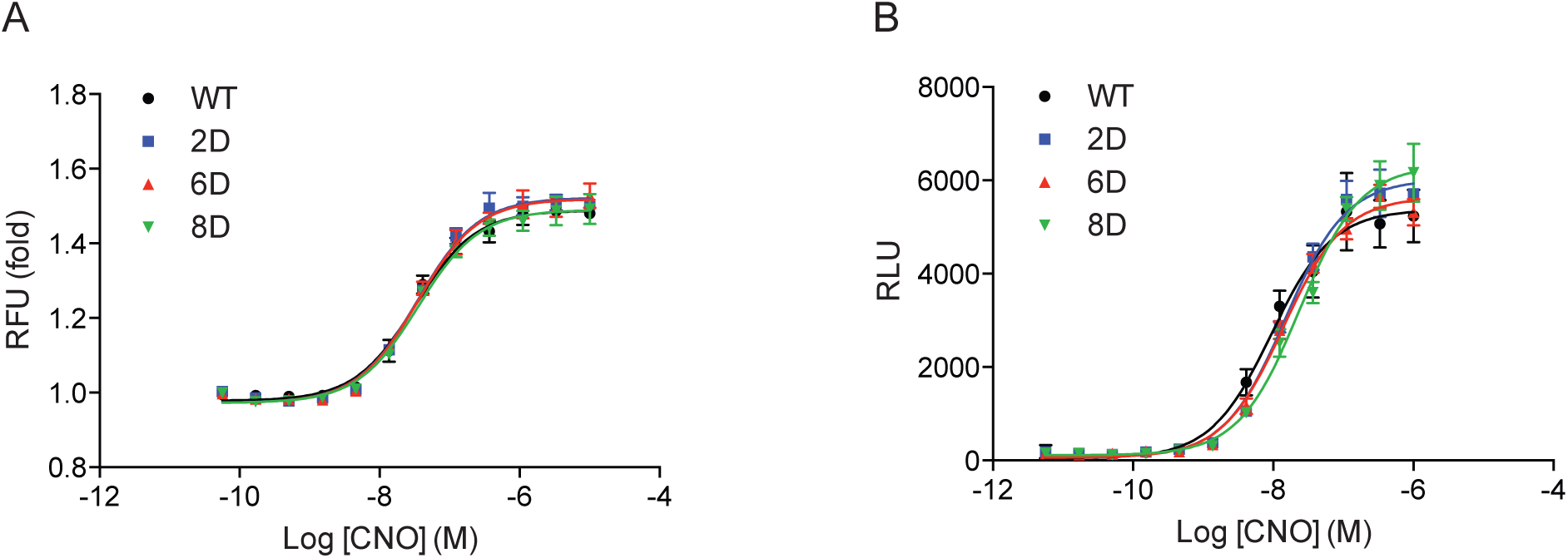
**(a)** FLIPR assay for validation of Gq activity of the phosphomimetic fusion receptor mutants. HEK293T cells transfected with hM3Dq-V2 tail-ABIcs (black line), hM3Dq-V2 tail (2D)-ABIcs (blue line), hM3Dq-V2 tail (6D)-ABIcs (red line), or hM3Dq-V2 tail (8D)-ABIcs (green line), were stimulated with CNO, and the Gq signal was measured by FLIPR^Tetra^. (b) SRE Luciferase assay validation of activity of the phosphomimetic fusion receptor mutants. HEK293T cells were co-transfected with SRE-Luc, PYLcs-β-arrestin2-YFP, and hM3Dq fusion proteins as follows; hM3Dq-V2 tail-ABIcs (black line), hM3Dq-V2 tail (2D)-ABIcs (blue line), hM3Dq-V2 tail (6D)-ABIcs (red line), or hM3Dq-V2 tail (8D)-ABIcs (green line). The cells were stimulated with CNO for 4 h, and then the luciferase activity was measured.

**Supplementary Figure 2.**
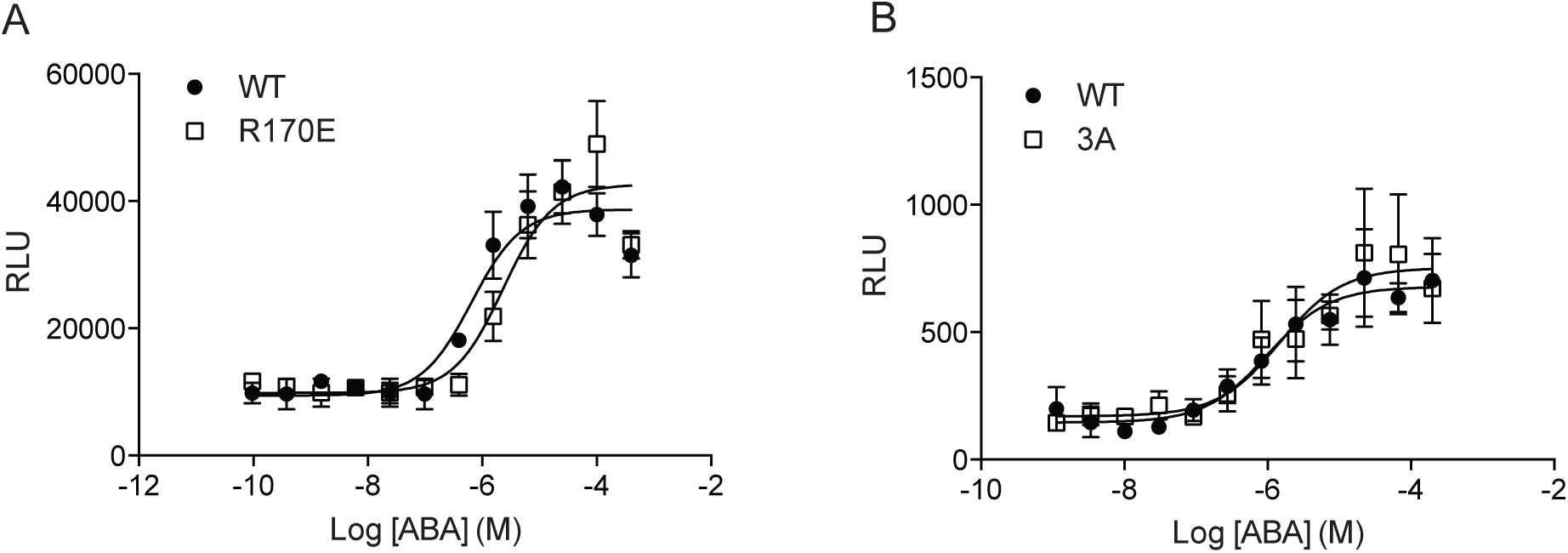
SRE Luciferase assay. HEK293T cells were co-transfected with SRE-Luc, hM3Dq-V2 tail-ABIcs, and PYLcs-β-arrestin2-YFP mutants as follows; (a) PYLcs-β-arrestin2-YFP (black line) or PYLcs-β-arrestin2 (R170E)-YFP (red line), (b) PYLcs-β-arrestin2-YFP (black line) or PYLcs-β-arrestin2 (3A)-YFP (red line). The cells were stimulated with ABA for 4 h, and then the luciferase activity was measured.

